# Machine Learning-Guided Antibody Engineering That Leverages Domain Knowledge To Overcome The Small Data Problem

**DOI:** 10.1101/2023.06.02.543458

**Authors:** Thomas Clark, Vidya Subramanian, Akila Jayaraman, Emmett Fitzpatrick, Ranjani Gopal, Niharika Pentakota, Troy Rurak, Shweta Anand, Alexander Viglione, Rahul Raman, Kannan Tharakaraman, Ram Sasisekharan

**Affiliations:** Altus Enterprises, 900 Middlesex Turnpike, Billerica, MA; Department of Biological Engineering, Koch Institute for Integrative Cancer Research, Massachusetts Institute of Technology, 77 Massachusetts Avenue, Cambridge MA 02139

**Author notes:** To whom correspondence must be addressed. Contributed Equally (Thomas Clark, Vidya Subramanian, Akila Jayaraman).

## Abstract

The application of Machine Learning (ML) tools to engineer novel antibodies having predictable functional properties is gaining prominence. Herein, we present a platform that employs an ML-guided optimization of the complementarity-determining region (CDR) together with a CDR framework (FR) shuffling method to engineer affinity-enhanced and clinically developable monoclonal antibodies (mAbs) from a limited experimental screen space (order of 10^2 designs) using only two experimental iterations. Although high-complexity deep learning models like graph neural networks (GNNs) and large language models (LLMs) have shown success on protein folding with large dataset sizes, the small and biased nature of the publicly available antibody-antigen interaction datasets is not sufficient to capture the diversity of mutations virtually screened using these models in an affinity enhancement campaign. To address this key gap, we introduced inductive biases learned from extensive domain knowledge on protein-protein interactions through feature engineering and selected model hyper parameters to reduce overfitting of the limited interaction datasets. Notably we show that this platform performs better than GNNs and LLMs on an in-house validation dataset that is enriched in diverse CDR mutations that go beyond alanine-scanning. To illustrate the broad applicability of this platform, we successfully solved a challenging problem of redesigning two different anti-SARS-COV-2 mAbs to enhance affinity (up to 2 orders of magnitude) and neutralizing potency against the dynamically evolving SARS-COV-2 Omicron variants.

## Introduction

Fueled by the success of AlphaFold, there has been tremendous interest in leveraging Machine Learning (ML) to engineer proteins, including antibodies for developing them as successful therapeutics in the clinic ^1^. ML-guided optimization of the target antigen-binding affinity of antibodies has enormous potential to generate novel antibodies with enhanced affinity and potency. A variety of deep learning (DL) models have been trained on datasets of experimentally generated libraries of antibody complementarity determining regions (CDRs). These DL models have been used to generate ‘optimized’ candidates faster than typical screening using display technologies ^2 2 3 4 5 6^. However, 10^4^-10^8^ designs are generally needed, thereby requiring a large experimental screening space, to analyze and verify the predictions from these models.

Unlike large antibody sequence datasets, the publicly available data that combines structural information with the experimentally measured effect of CDR mutations on antibody-antigen affinity (for example the SKEMPIV2 database) is limited. Moreover, the datasets in the SKEMPI database are biased towards mutation of CDR residues to alanine. The alanine mutation is a restricted subset relative to those sampled during *in vivo* affinity maturation or by an expert with extensive domain knowledge on antibody-antigen interactions. DL models including GNNs pre-trained on PDB ^7^ or on SKEMPIV2 ^8^ have achieved modest *in silico* benchmarks for successfully predicting affinity enhancing mutations despite the potential overfitting of the model parameters on limited datasets. In the case of ^8^, the model predicted-mutations were validated by experimental screening of less than 100 mutations for their impact on neutralization of a SARS-COV-2 antibody. However, given the potential bias in the alanine mutation training dataset, the robust performance of these models to consistently optimize affinity across different antibodies using a small experimental screen size remains to be evaluated.

With public antibody repositories such as Observed Antibody Space (OAS) ^9^ that contain next generation sequencing (NGS) data growing to billions of sequences, it portends the significance to generate diverse, naturally occurring, antibodies using large language models (LLMs). An ensemble of pretrained LLMs were also recently used to predict affinity enhancing mutations on a variety of anti-viral antibodies in a structure and antigen independent manner, but using these LLMS did not uniformly improve binding, including in the context of the antibody binding to SARS-COV-2 Omicron variant ^10^. Therefore, there are challenges with using pre-trained LLMs to predict affinity enhancement as these models do not consider the protein-protein interaction data which fundamentally limits the design space.

The recent emergence of employing DL models in antibody discovery has led to an overwhelming trend of abandoning human engineered features for the hopes of implementing an end-to-end machine-guided *de novo* antibody design. While this trend has gained popularity, there are still gaps and challenges in using DL models for practical or translational antibody engineering. Our approach over the last 10 years has relied on engineering *in silico* metrics (or features) using network-based representations of protein structure and protein-protein interactions. We have used these features to guide human selection of affinity enhancing mutations ^11 12 13 14^. Using an iterative design cycle of computational generation of a small screen size (fewer than 100 constructs per cycle) and experimental evaluation (wet-lab screening of expression, yield, purity, and target binding affinity), we have successfully engineered diverse clinical-stage antibodies with specificity and enhanced affinity to a target epitope on various antigens ^12 13 15 16^. Notably, even within the small screen size, our overall approach permitted us to sample diversity in the sequence and structural features that loop-back into our design platform for learning.

With this foundation, we have developed an ML-guided antibody engineering platform that combines knowledge-guided featurization with a data-driven model design. Notably, the ML model presented here, referred to henceforth as the Antibody Random Forest Classifier or AbRFC, outperformed the GNN and LLM-based models on an in-house validation dataset comprising diverse CDR mutations for affinity enhancement, thereby representing an important step towards addressing the overfitting of the limited antibody-antigen interaction data. Also, unlike other ML-models that used regression to optimize mutations for affinity enhancement, the approach described here uses classification, classifying deleterious from non-deleterious mutations to maximally leverage the information content in alanine mutations (ALA-scan) heavy SKEMPIV2. Importantly, the individually classified affinity-enhancing mutations on different CDRs were combined to generate synergistic enhancement of antigen-binding affinity. The optimized CDRs were then shuffled with FR regions that were selected from the large antibody sequence repositories to generate candidate antibodies that were simultaneously optimized for antigen-binding affinity and developability.

Using the ML-guided platform described here, we engineered two distinct antibodies respectively building on two starting template antibodies that were developed against the original Wuhan SARS-COV-2 virus and that lost potency against Omicron and subsequent variants. For each antibody, the CDR optimization and CDR-FR shuffling each involved experimental screen size of fewer than 100 constructs. The engineered antibodies showed up to two orders of magnitude improved affinity compared to the corresponding template mAbs against various Omicron subvariants BA.1, BA.2 and the more recent VOCs BA.4/5, and XBB.1, surpassing the results produced by a pretrained LLM on S309 ^10^. Significantly, using a combination of these two mAbs shows potency (IC_50_ < 200 ng/ml) against the different Omicron subvariants.

The ML-guided platform described in this study highlights the value of employing classical ML methods (e.g. linear regression, support vector machines, and tree-based boosting or bagging methods) with expert-guided feature engineering and an appropriate choice of training and validation datasets in achieving affinity enhancement with only two rounds of experimental screening of order of 10^2^ constructs.

## Results

ML-guided antibody engineering platform described in this study is built over the foundational concept of “lab-in-a-loop” (**Figure 1A**) that we have evolved over the past several years. This platform involves two iterative cycles, the *first* for optimizing CDRs to enhance binding affinity and the *second* to select appropriate scaffolds to combine with the optimized CDRs to generate lead candidates. The candidate sequences in the first iteration were generated by *in silico* saturation mutagenesis of the template antibodies’ CDRs. These *in silico*-generated candidates were virtually screened using the AbRFC model to predict non-deleterious mutations (since the model is set up classify deleterious vs non-deleterious mutations). The rationale for predicting the non-deleterious mutations was that experimental screening less than 100 candidate mutations (e.g. “selected subset” in **Figure 1B**) on a 96-well plate will potentially result in affinity enhancing mutations in a single round. The second iteration starts with search for antibody scaffolds in the large antibody sequence databases (including OAS) to select appropriate FR regions that can be combined with the optimized CDRs from the first iteration as previously described ^16^ to generate sequences with various FR-CDR combinations. These second-round candidates were virtually screened to select candidates that have optimal developability scores (sequence liabilities, charge distribution, hydrophobic patches, T-cell epitopes, etc.) that are computed *in silico*. The final candidates were experimentally tested for critical quality attributes including potency.

**Figure 1.**
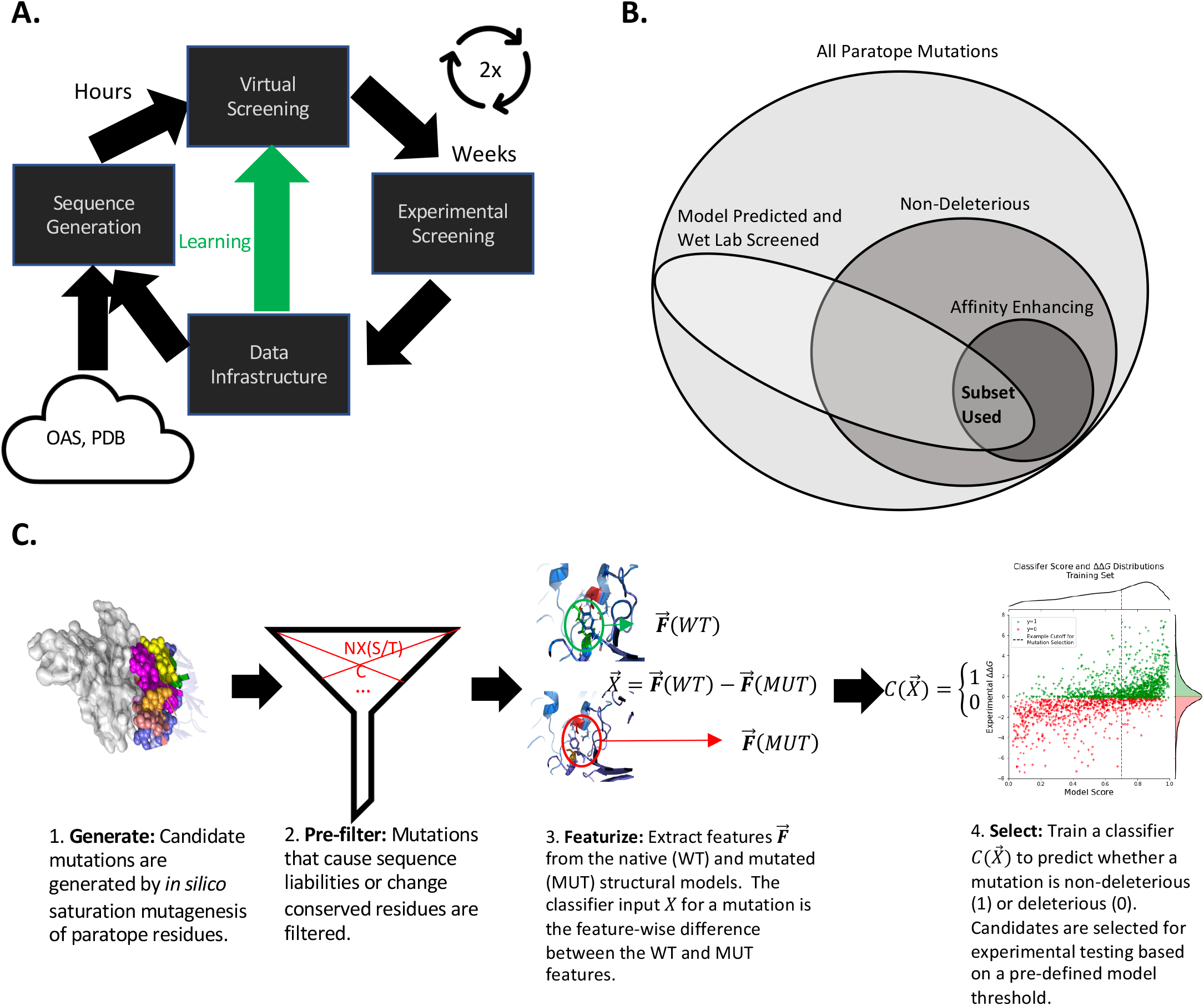
Schematic of our ML-guided Platform. **A**. The underlying lab-in-the-loop screening methodology is enabled by 4 key pieces. First, an *in silico* algorithm that uses large public datasets and in-house data to generate candidate sequences. Second, a virtual screening step, where the sequences are screened for properties such as affinity and developability. Finally, constructs are screened in the wet lab and data is stored so that it can be used both for generating subsequent constructs and for enhancing the virtual screening models. The designs for the G and C programs required only 2 iterations of this cycle. **B**. The filtering hypothesis that is central to our novel approach posits that a model that is trained to predict the binary *deleterious* vs *non-deleterious* classification task will filter the candidate set sufficiently such that affinity enhancing mutations will be observed during wet lab screening. **C**. The *in silico* procedure used to identify candidate mutations in practice. Steps 1-3 are typical for *in silico* affinity prediction pipelines, with many different featurization techniques used. Our approach is differentiated by splitting the training dataset into binary class labels rather than training a model to predict the experimental **ΔΔ***G* directly and using a cutoff ΔΔ*G* slightly below 0 to account for noise in the experimental measurements.

### The AbRFC model for ML-Guided Affinity Optimization

The common components of building any ML model are the objective function, the model architecture, and featurization (**Figure 1C**). The model design choices depend on the size, type, and quality of the training dataset. Despite the wealth of antibody sequences, only limited data on antigen-binding affinity including impact of CDR mutations on binding is publicly available. A subset of the publicly available protein-protein binding affinity database SKEMPIV2 ^17^ containing point mutations on antibody-antigen complexes (“training dataset”) is relatively small (N=900) and overwhelmingly biased (**Figure 2A**) (61% of the mutations are to ALA). In contrast to the training dataset, the validation dataset included in-house data (unpublished) from a different antibody-antigen affinity enhancement campaign, wherein the point mutations were predicted by an expert using a structural model of antibody-antigen complex and were validated for binding experimentally. Therefore, the validation dataset contained very few mutations to ALA (**Figure 2B**), and is enriched in patterns that the expert has learned to apply in a residue-specific manner – for example in this instance frequently mutating L to F and Y and S to R and K. This led us to reason that *any* model trained on the available limited training dataset would be biased relative to a model trained to learn affinity enhancement specifically, but that the training dataset still contains relevant information.

**Figure 2.**
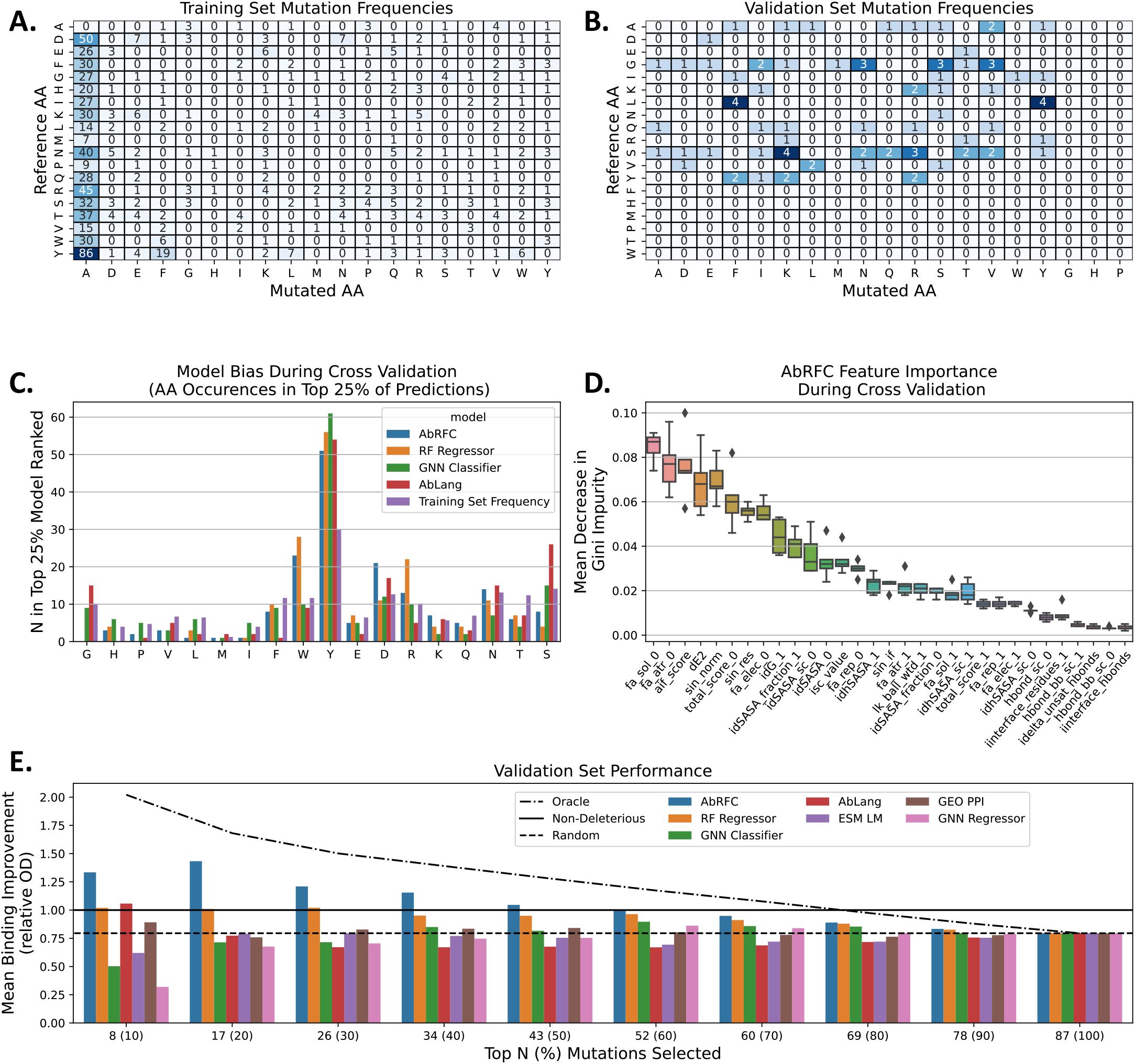
Dataset Analysis for Model Design Decisions. **A**,**B**. Distribution of sampled mutations in **A**. the training subset of SKEMPIV2, and **B**. The validation set, which includes expert-picked mutations for an affinity enhancement campaign. The distribution of mutations that are picked during an affinity enhancement campaign is fundamentally different from the ALA scan heavy dataset that is publicly available for training. A ML model will need to generalize from the ALA-scan heavy training dataset to the affinity enhancement use case. **C**. Different models learn different biases, despite being trained and evaluated on identical CV splits. **D**. Feature importance calculated for the AbRFC during cross validation show that the strongest predictors include both local (fa_sol_0,fa_atr_0, aif_score) and global descriptors (dE2,sin_norm). **E**. We ranked the mutations in the IH dataset generated for expert-picked mutations using each of the models and plotted the mean relative OD of the top N ranked constructs. The Oracle line shows what a perfect ranking would look like, while the random line indicates the performance of random reranking. AbRFC performs the best on this (N=87) validation set, while the AbLang language model also selects non-deleterious mutations in the top 10%.

To mitigate these challenges, we made three model design choices. First, we chose to frame the prediction problem as a binary classification problem rather than using the typical regression formulation. Classification is a more natural fit for the filtering objective of virtual screening and also reduces the influence of outliers (e.g. mutations with reported fold change of >10000, |ΔΔ*G*| > 5.45) on the model predictions. Additionally, rather than trying to predict affinity-enhancing mutations (|ΔΔ*G*| ≫ 0), our model predicts deleterious vs. non-deleterious mutations (|ΔΔ*G*| > −*ϵ*). We reasoned this formulation is more likely to learn generalizable patterns from an ALA-scan heavy training dataset and chose a small offset ϵ from |ΔΔ*G*| = 0 to avoid assigning arbitrary class labels due to experimental noise.

Second, we used a classical ML algorithm with engineered features rather than a DL approach. Specifically, we selected a Random Forest Classifier where the bootstrap aggregating (bagging) nature of the algorithm naturally reduces model variance (errors due to small changes in the training dataset) particularly for large forest sizes ^18^. This method is also amenable to additional regularization by changing hyper parameters, which we leveraged, choosing a minimum of 10 samples per leaf and a large forest size of 1000 trees.

Third, we employed a defined set of features of antibody-antigen interactions that were specified by human experts. These expert-engineered features, in contrast to the high-dimensional representations learned by DL algorithms, have the advantage of encoding previously validated inductive biases that are less likely to be overfit to the publicly available small, and ALA scan-biased training dataset. In addition to a variety of Rosetta energetics terms accessed in PyRosetta ^19^, we used previously validated statistical ^12^ and network-based ^11^ features. The Random Forest Classifier applied to this featurization is termed AbRFC.

### Training dataset Cross Validation

To assess impact of our modelling choices on the performance during cross-validation on the training dataset, we compared our model to a retrained classifier using the architecture from ^8^ (GNN Regressor), a RandomForestRegressor (RF Regressor) with the same parameters as AbRFC, and an LLM-based model that predicts mutations in a similar manner to Hie BL et. al., ^10^, but using an antibody-specific LLM (ABLang) ^20^. All models performed similarly during cross-validation (**Figure S1**), except that the LLM was slightly worse on some folds (still surprisingly good given it uses no antigen information), which was expected since all the folds in the training dataset contain a large amount of ALA-scanning data. However, most mutations encountered during virtual screening (18/19 for *in* silico saturation mutagenesis) are not ALA mutations.

Notably, the cross-validation showed that model architecture and objective function influence AA mutation preference, even when using the identical training dataset. Both RF models prefer charged residues, whereas the GNN Classifier predicts backbone altering G and P mutations at approximately the underlying training dataset frequency, unlike the RF algorithms that almost never predict these changes. The bias of our featurization against G and P arises from our network and statistical metrics not accounting for the potential backbone altering effects of substituting to these amino acids. All models are significantly enhanced for aromatic residues, but ABLang and the GNN classifier have a stronger relative preference for Y, while the RF algorithms have increased W preference compared to baseline (**Figure 2C**). This indicates that when scoring mutations in practice there will likely be bias due to the underlying design choices.

Feature importance scores generated during cross validation show the following top 5 features for differentiating non-deleterious and deleterious mutations – two Rosetta energy terms (fa_sol_0 and fa_atr_0), a statistical amino acid preference feature ^12^, a networking feature of Ab-Ag complex (sin_norm), and a total energy difference (dE2) (**Figure 2B). Table S1** details the features used in the AbRFC.

### Testing against the Validation Dataset

The validation dataset consists of mutations selected by an expert to enhance affinity and therefore it has its own bias and is not representative of saturation mutagenesis. We used this dataset to test our ML model and others including GNN Regressor, GeoPPI ^7^, and an ensemble of ESM Language models ^21^. AbRFC picked (on average) non-deleterious mutations in the top 10%, 20%, 30%, and 40% of ranked mutations **(Figure 2E)**. AbLang was the only other model able to predict non-deleterious mutations in the top 10% of ranked mutations on the validation dataset. To ascertain whether our model would generalize to the saturation mutagenesis setting, we applied it to engineer two antibodies targeting two distinct epitope regions on SARS-COV-2 with the goal of providing a long-lasting solution to the dynamically evolving SARS-COV-2 virus.

### Starting template antibodies targeting mutationally constrained epitopes on SARS-COV-2

An important part of the platform is to engineer antibodies targeting specific epitopes. In the context of SARS-COV-2, our goal was to identify epitopes that are constrained to mutate. To this end, we ranked template epitopes using a metric that combines the sequence conservation of epitope residues with their networking to CDR residues (**Supplementary Methods**). Among the various epitope regions, those that were targeted by a class III mAb (S309) ^22^ referred to here as ‘glycan’ epitope (since it comprises of glycosylation site) and class IV mAb CR3022 ^23^ referred to here as ‘cryptic’ epitope (since it is harder to access on the spike rimer) were highly conserved across SARS-COV-1 and SARS-COV-2 (**Figure S2**). Therefore, we chose CMAB0 (a CR3022 derivative) and GMAB0 (S309) as starting template antibodies to engineer new antibodies with substantially enhanced affinities to the Omicron variant and subvariants given that CMAB0 and GMAB0 had significantly lower binding affinity to these variants.

### First round of experimental screening: screening for affinity enhanced CDRs against BA.1

Two crystal structures per template (7BEP/7R6W for GMAB0, 6YLA and 7LOP for CMAB0) were used to minimize structure-based variability. The BA.1 epitope mutations (G339D/N440K for GMAB0 and S371F/S375F for CMAB0) were modeled using PyRosetta (see **Methods**). We then ranked mutations based on the AbRFC model score (predicted non-deleterious class probability). We noticed that ranking mutations purely based on model score without considering other factors naturally oversamples specific positions and mutant AAs (**Figure 2C**). As stated previously our model design was such that it has a bias against mutating to glycine and proline residues. However, glycine mutations have been known to contribute to affinity and specificity ^24^ and were preferred by other models (**Figure 2C**). Therefore, to address these biases and enhance diversity in the selected candidate set we added additional rules to augment the AbRFC score with Rosetta ΔΔG and ABLang Score (unbiased by training dataset) when considering mutations to Glycine (see **Methods**).

An ELISA-based screening assay using the BA.1 RBD (see **Methods**) showed that 21 and 24 of the ML-filtered CDR mutation sets had improved binding to the glycan and cryptic epitopes over GMAB0 and CMAB0 respectively (**Figure 3 A,B,C,D)**. Included in this set are several mutations such as H31_SK,H100_AS,L52_SE for the glycan antibody and H55_SQ, H99_SN/W, L31_NE, L90_QN and L93_SK for the cryptic antibody that have significantly higher OD values. This shows filtering for non-deleterious mutations and using diverse sampling is sufficient to arrive at multiple potential affinity enhancing mutations. In the next step, we combined CDR loops carrying the affinity enhancing mutations with diverse FR regions to derive affinity-enhanced, developable candidates.

**Figure 3.**
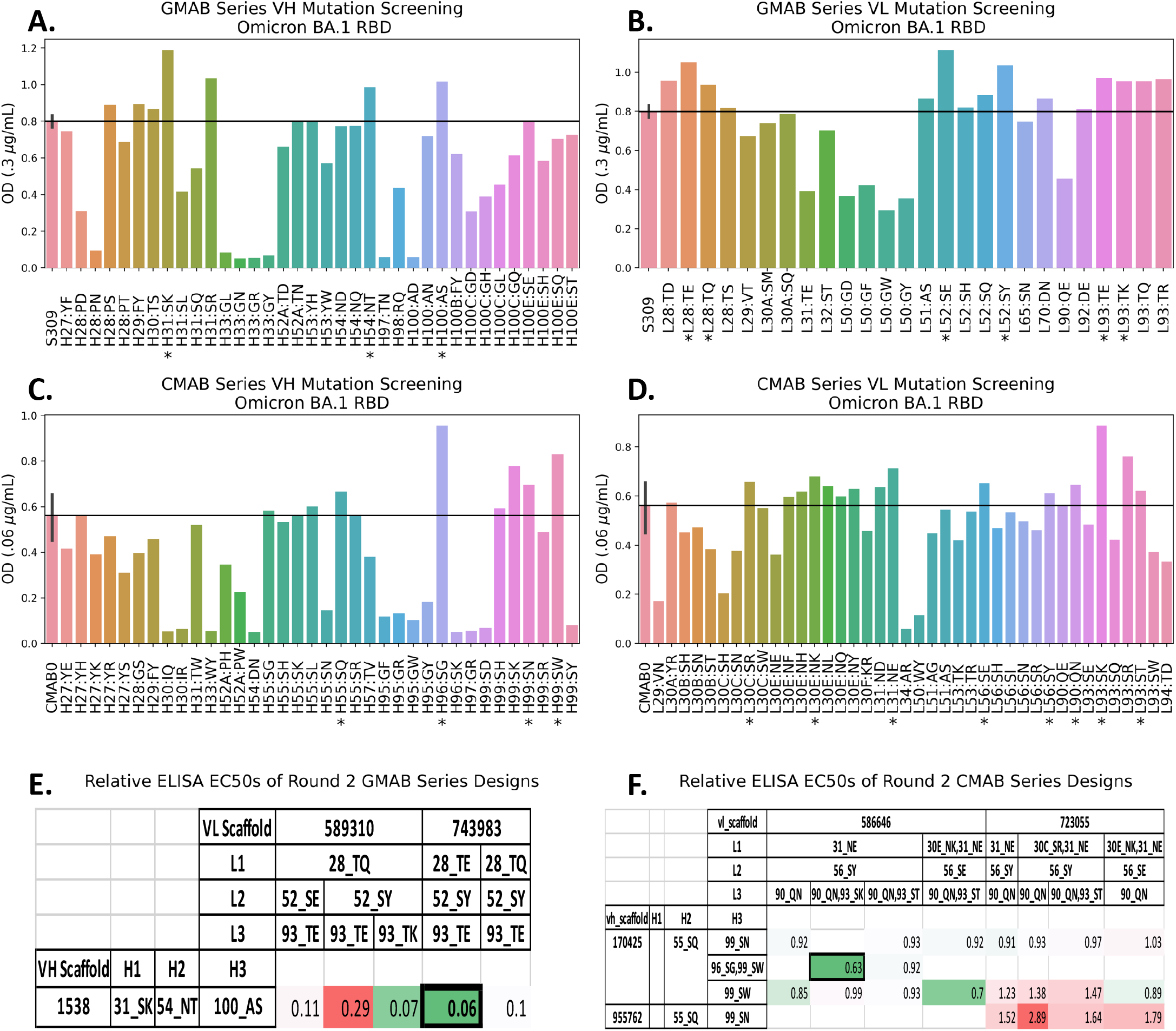
Experimental Screening of Point Mutations and CDR-FR Shuffling. **A**. Single concentration ELISA OD to estimate the binding of constructs carrying the S309 VH point mutations to the BA.1 RBD (* indicates point mutation used in round 2). **B**. Single concentration ELISA OD to estimate the binding of constructs carrying the S309 VL point mutations to the BA.1 RBD. **C**. Single concentration ELISA OD to estimate the binding of constructs carrying the CMAB0 VH point mutations to the BA.1 RBD. **D**. Single concentration ELISA OD to estimate the binding of constructs carrying the CMAB0 LH point mutations to the BA.1 RBD. **E**. Relative EC50 (relative to S309) of glycan epitope targeting antibodies constructed using the selected point mutations and FW-CDR shuffling. Rows indicate H FR and HCDRs used, while columns represent the VL FRs and CDRs. Details on the VH and VL FRs are in **Table S2**. The most promising antibody (red box) was selected for further characterization. **F**. Relative EC50 (relative to CMAB0) of cryptic epitope targeting antibody designs. The most promising antibody (red box) was selected for further characterization.

### “Lab-in-a-loop” second iteration: CDR-FR shuffling to further optimize the affinity enhanced candidates

Given that CDRs and FR regions play a key role in antibody affinity, stability and pharmacokinetic properties ^25^, the second round of designs combined the top point mutations with novel human VH/VL frameworks to make clinically developable lead candidates. This iteration was critical to address issues of affinity-enhancing CDR point mutations could potentially introducing undesirable physical properties (alter expression, Fab melting temperature (Tm), isoelectric point (pI) as observed in other studies ^25 26^. Importantly, we directly combined the promising affinity enhancing mutations rather than building mutants up gradually through multiple rounds, permitting us to generate lead candidates in 2 iterations.

A diverse set of FR regions were sampled from a database containing next-generation sequences from public repositories such as cAb-Rep ^27^ and OAS. As previously described ^16^, FRs were filtered based on structural properties such as North CDR cluster ^28^, CDR length (to accommodate the modified CDR loops), PTMs, rare amino acids, high-energy (PyRosetta) residues, and FR sequence diversity. **Figure S3** shows the AbLang embeddings of the scaffold sequences, where the spread of the red crosses illustrates the diverse sequence space sampled.

Second round experimental screening significantly showed affinity enhanced constructs with developability properties that are in the range seen for clinical stage therapeutic antibodies, both by *in silico* (**Figure S4**) and experimental (**Table S3**) analyses. The combination of scaffolds and point mutations that were screened by ELISA against BA.1, along with the relative EC50 of the designs with respect to the template (**Figure 3 C,F**). The most promising of these leads were then tested in Octet for binding against the BA.1, BA.2, BA.4/5, and XBB.1 subvariants. The lead candidate targeting cryptic epitope, CMAB283, showed significantly increased affinity to BA.1 (to which the affinity was optimized), but also to the other VOCs, BA.2, BA.4/5, BQ1.1, and XBB.1, maintaining a KD of 2.14E-11 or lower on all tested VOCs relative to >1nM KD for the template CMAB0 (**Figure 4A, Figure S5**). The lead targeting glycan epitope, GMAB156 showed improved affinity ranging from ∼317-fold for BA.1 to 1.2-fold for XBB.1. Importantly, GMAB156 maintained sub nanomolar binding on all VOCs while S309 exhibited showed KDs >1 nM for BA.2 and, BA.4/5 (**Figure 4A, Figure S5**). Given the established correlation between affinity and neutralization for S309 ^29^, the consistent sub-nanomolar binding of GMAB156 makes it an attractive therapeutic candidate, especially since the current circulating strain (XBB.1.5) has no mutations near the glycan epitope relative to XBB.1. Given that our lead mAbs showed substantial improvement in the binding affinity, we tried to rationalize the positive effect of these mutations on the antigen-binding use three-dimensional structural models of our lead mAbs with the various SARS-COV-2 variants RBD (**Supplementary Methods**). However, while some of the mutations selected by the platform were rationalizable using structure-based reasoning, the platform also discovered mutations that are not obvious *a priori* from static structure analysis (**Supplementary Information**).

**Figure 4.**
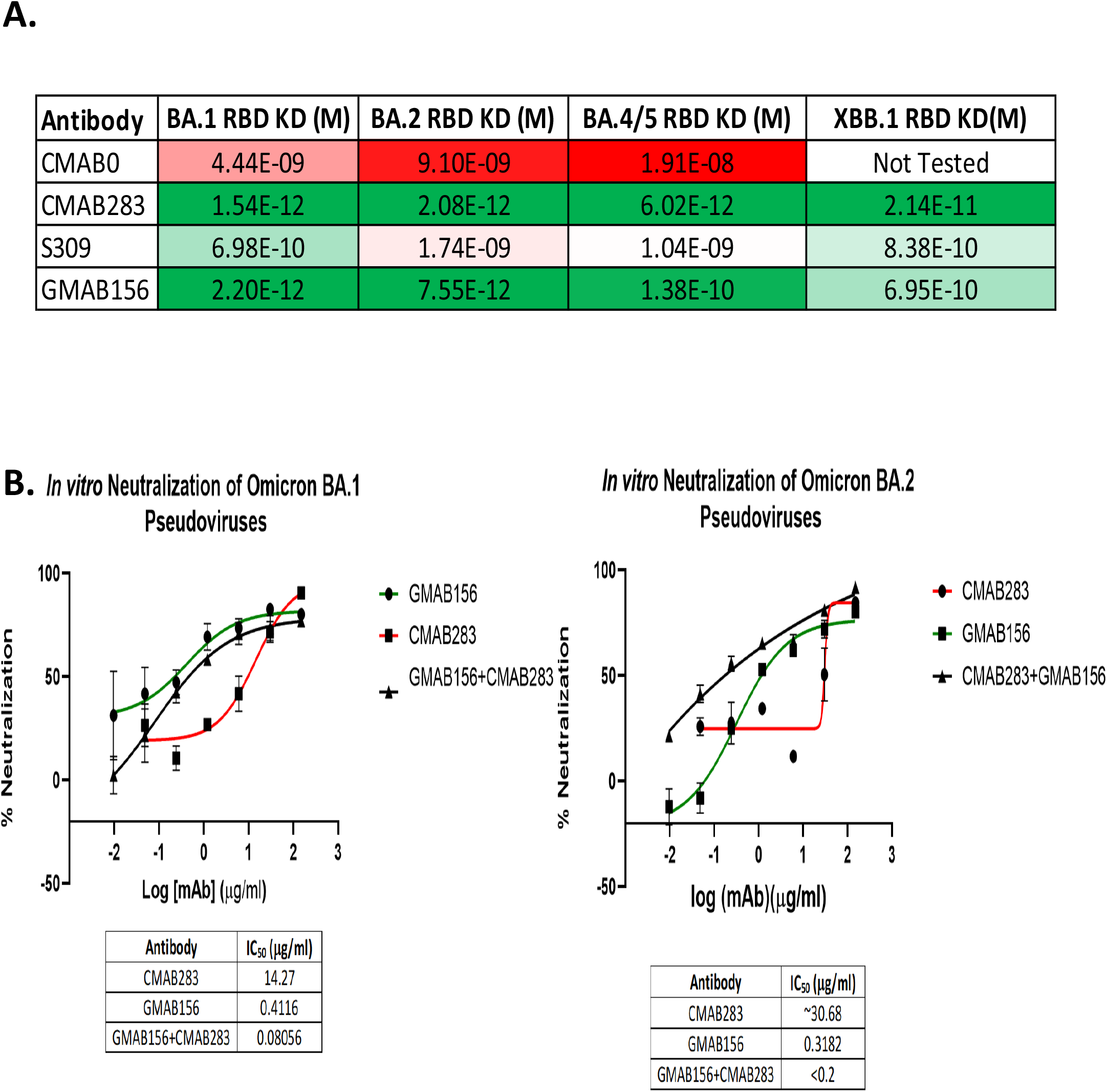
Binding affinity and pseudovirus neutralization of the engineered leads. **A**. K_D_ values of the templates and engineered leads when measured by BLI. The engineered leads show significantly (7-1000x) improved affinity relative to the templates across VOCs. **B and C**. The neutralization profile of the antibodies against the designed strain (BA.1) and the subsequent strain (BA.2) is consistent with the enhanced affinity and there appears to be some synergistic effect from targeting the orthogonal epitopes.

### Pseudovirus Neutralization of SARS-COV-2 variants by the lead mAbs

The two candidate mAbs GMAB156 and CMAB283 that showed improved affinity (as compared to the respective template mAbs) to the distinct glycan and cryptic epitope regions on Omicron subvariants BA.1, BA.2, BA.4/5, and XBB.1 respectively were tested further. Especially, we evaluated the impact of improved binding affinity and the postulated synergy (assessed using our previously described PADS framework) ^30^ in targeting multiple epitopes on the neutralization potency. We adapted a SARS-COV-2 pseudotyped virus neutralization assay based on previously established methods ^31^. As reported previously, the neutralization of the template mAb S309 significantly reduced between Omicron and subsequent BA.2 and newer VOCs including BA.4/5 and others ^31^.The loss of neutralization potency of S309 correlates with its binding affinity to the corresponding spike proteins of the Omicron subvariants ^29^ (**Figure 4A**). On the other hand, our candidate GMAB156 shows uniformly potent neutralization across BA.1 and BA.2 which correlates with its substantially higher binding affinity to the corresponding spike protein (**Figure 4B and 4C**). Given this correlation and based on its sub nanomolar binding to BA.4/5, BQ1.1, and XBB.1, we anticipate that this mAb would show substantially improved potency to these more recent subvariants to be effective as a therapeutic to counter these subvariants.

Unlike the glycan epitope, substantial improvement in the affinity of the CMAB283 to the cryptic epitope when compared to CMAB0 does not lead to potent neutralization of BA.1 and BA.2 Omicron variants. This is likely due to the ‘cryptic’ nature of the epitope that might have different exposure in the RBD of spike protein when compared to the spike protein on the surface of the pseudovirus or live virus. However, the combination of CMAB235 with GMAB156 further enhances potency of GMAB156 both against BA.1 and BA.2 (**Figure 4B and 4C**) indicating synergy in targeting these antigenically distant epitope surfaces on the spike protein. Additionally, using a combination therapy is attractive as it minimizes the chances of mutational escape.

## Discussion

*In silico* antibody design is a rapidly evolving field that leverages the power of computational methods and molecular modeling to create epitope-specific antibodies with specific properties. While *de novo* design of epitope-specific antibodies is still a challenge, optimization of antibodies known to target the desired target epitopes is possible using structure-guided metrices and ML algorithms.

The earliest efforts to introduce ML in computational antibody engineering was made by us where we combined feature engineering and logistic regression to develop a predictive model for discriminating native antigen-antibody poses from decoys ^12^. Classical ML models can work on smaller (training) datasets and are computationally cheap and readily interpretable. On the other hand, deep learning models require extremely large datasets to train and are difficult to interpret owing to their “black box” nature. While tools like mCSM-AB2 ^32^ have employed classical ML to determine the impact of mutations on antibody binding, this represents the first study (to the best of our knowledge) to show that antibodies having improved properties (e.g., affinity) can be generated within a small experimental screen space using classical ML. The classical ML-guided platform presented here is distinct in the following aspects. We reduce the challenge of predicting the most affinity enhancing mutations to a classification problem, where we train a model to discriminate deleterious mutations from neutral or affinity enhancing mutations in the CDRs. Consequently, using the classification models, we were able to sample diverse mutations and rapidly screen combinations of mutations experimentally despite the size and bias of the training dataset. Additionally, we have explicitly examined the biases of our model and those implemented by others and accordingly augmented the AbRFC model scores to sample diverse mutations including glycines. Minimizing these biases and increasing the mutation sampling led to increased propensity for identifying affinity enhancing mutations within a small screen size. A two-pronged approach of CDR engineering (using ML-guided mutations) and scaffold selection was employed to identify antibody FR regions from large sequence databases that have improved properties, which translate to better developability criteria as a clinical stage therapeutic. Finally, another important distinction of the platform is the richness of the datasets on single and combination point mutations on the CDRs and their associated experimental binding and developability metrices. The progressive increase in these datasets further augment the classification performance of the model for future tasks thereby making the ML-guided lab-in-a-loop more efficient with each antibody engineering campaign.

Acknowledging the limitations in the relatively small and ALA-scan heavy publicly available training datasets, we set out to apply ML specifically to pre-screen a large set of point mutations. Recent work on activity cliffs in small molecules ^33^, an analogous situation where very small differences can have large impact, supports the idea of the continued use of classical ML for the problems involving predicting impact of small molecular changes on overall function and other properties, despite the exciting capacity of deep learning to generate large sets of highly diverse sequences

To ensure diversity in our screening, we used basic rules including structural location and physiochemical properties in the context of CDR engineering and ported the CDRs onto a diverse set of scaffolds that passed *in silico* developability filters. One can possibly imagine using a more sophisticated system such as generative modeling to sample from the distribution of somatic hypermutations associated with a template clonotype and subsequently scoring them with this platform to achieve a fully end-to-end AI system. Further, these models may be able to suggest and score candidates that have multiple mutations, decreasing the experimental iteration time still further.

The platform presented here demonstrates that given a template antibody and structural information of the antibody in complex with a homologous but nonidentical antigen, it is possible to apply the ML-guided computational methods presented here to find highly diverse, functionally improved antibodies using only 2 rounds of experimental iterations. In conclusion, we have successfully applied this platform to a practical problem of obtaining rapid antibody neutralization solutions to the constantly evolving SARS-COV-2 viruses.

## Methods

### Training Dataset

The GNN and Random Forest models were trained on the mCSM-AB2 dataset (http://biosig.unimelb.edu.au/mcsm_ab2/data). AbLang is pretrained and therefore was not retrained.

### Structure Processing

All structures, regardless of dataset, were processed identically. PDBs associated with the complexes in the training dataset were downloaded from the RCSB Protein Data Bank. Structures were cleaned by renaming heavy and light chains to (H,L respectively) and renumbering them using the Chothia numbering scheme. Structures associated with the original (WT) complex were relaxed 10 times using Rosetta Fast Relax with the identical parameters to those used in the RosettaAntibodyDesign protocol. The lowest energy structure was selected and used for feature extraction. For each mutation, a new PDB file was generated using PyRosetta, wherein side chains within 5A heavy atom distance of the mutation were repacked.

### Classification Labels

To extract signal from the available training dataset, a cutoff of fold 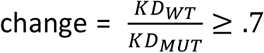 was used to classify mutations as non-deleterious (*y* = 1) or deleterious (*y* = 0). ΔΔG = ΔG_wt_ − ΔG_mut_ values were converted to fold change using:

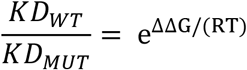

The cutoff of .7 was used rather than 1 to reduce the signal noise for mutations with .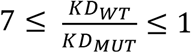.

### AbRFC Implementation

The Random Forest algorithm was implemented using the RandomForestClassifier from the scikit-learn package in python. A small number of examples in the training dataset are not AB-AG complexes and therefore do not permit the use of the AIF score. AIF scores for these instances were imputed using the IterativeImputer class from scikit-learn.

Feature engineering was implemented by computing previously validated metrics for the mutant residue, the neighborhood residues (see **Supplementary Methods** for neighborhood definition), the interface residues (PyRosetta definition), or the full complex and used as inputs to the model. For each feature, the value of that feature used in the model is Δ*Feature* = *Feature*(*WT*) − *Feature*(*Mut*). Features ending in the suffix _0 are associated with the residue that was mutated, ending in the suffix _1 are neighbor features and otherwise are full complex or interface features. All features used are in **Figure 2D**.

Feature selection was performed by averaging the feature importance (calculated by the scikit-learn RandomForestClassifier based on the mean decrease in impurity) from all trees across all 5 CV folds.

The only hyperparameter that was tuned was the fold change cutoff value for the classifier. This was tuned by trying the values [.6,.7*,1,1.2], which resulted in spearmanr of [.53,.51*,.26,.002], indicating that predicting non-deleterious values resulted in better performance on the validation set. However, we note that the RandomForestClassifier still predicts non-deleterious mutations on average in the top 10% for all cutoff values tried: [1.33,1.42*,1.47,1.13], suggesting that the classification approach retains signal on the validation set regardless of cutoff.

### GNN Model Classifier Implementation

The code for the graph neural network was cloned from https://github.com/HeliXonProtein/binding-ddg-predictor. To retrain the network as a classifier rather than a regressor, a sigmoid layer was added to the final layer readout to map the embeddings to [0,1]. Rather than using a mean squared error loss, the cross-entropy loss was used. Hyperparameters for model training were extracted from the paper.

### AbLang Model Implementation

The AbLang package was installed via PyPi. The pre-trained models for the heavy and light chains were downloaded. The likelihoods for mutations at each position were scored using the ‘model(seqs,mode=‘likelihood’) command, and the logits were converted into probabilities using the softmax function.

### Cross Validation

The 1800 datapoints (representing 900 mutations) were split into 5 folds such that approximately 1450 datapoints were in the training dataset and ∼350 data points were in the validation set for each fold. Mutations from the same complex were always grouped into either the training or the validation set.

### Random Forest Regressor Implementation

The Random Forest Regressor was implemented using identical parameters except for those parameters that are classification specific (min_impurity_decrease, class_weight, criterion). The exact parameters used are in the **Supplementary Methods**.

### GNN Model Regressor Execution

The model was downloaded from https://github.com/HeliXonProtein/binding-ddg-predictor and the ‘predict.py’ script run as instructed in the README.md documentation. The same structures used to train AbRFC were input as the wild type and mutant pdbs for the predict.py script.

### GEOPPI Model Execution

GEOPPI was cloned from ‘https://github.com/Liuxg16/GeoPPI.git’ on 09/27/2021. Mutation candidates were evaluated using the command ‘ python run.py [pdb file] [Mutation] [partnerA_partnerB]’ as described in the README.md file.

### ESM Language Model Execution

The same ESM language models used in the github repository associated with Hie BL et. al.,^10^ were used to score the validation set mutations. Because the process used therein selects specific amino acids rather than scoring each mutation, we instead extracted a score for each mutation by computing 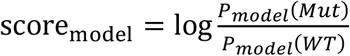.

### Structural Modelling of the Mutations

For S309, the PDBS 7BEP and 7R6W were used and for CMAB0 the PDBs 6YLA and 7LOP were used. The Omicron BA.1 mutations were performed using PyRosetta, with sidechain repacking with 5A of the mutated residues. For CMAB0, the mutations between CMAB0 from CR3022 were also added to the model prior to relaxation.

### Mutation Scoring

Residue positions to be mutated were selected by considering any position that had a heavy atom within 10A of the RBD in the starting structure (the loose criterion was to accommodate longer side chain mutations). Additionally, the following positions were ignored: template AA is Cysteine; template AA is Glycine AND is 90% conserved in human antibody sequences; template AA is any amino acid AND 95% conserved in human antibody sequences. All amino acids except Cysteine and AAs that would introduce glycosylation were considered. Mutations were ranked on a *position-specific* basis using the AbRFC model scores to ensure we could meet the following diversity requirements: <=7 mutations per position, >=1 non-synonymous mutation sampling in all CDRs. Additionally, as described above due to the bias of AbRFC against G mutations, decisions for Glycine mutations were made based on AbLang and Rosetta ΔΔ*G* scores in addition to AbRFC scores.

### CDR FR Shuffling

FRs were selected using an identical procedure to that described earlier ^16^. Briefly, filters for the S309-based designs included:

- H1 north cluster = H1-13-A, H2 north cluster = H2-10-A, H3 length = 18, maximum number of PTMS: 18, maximum rare amino acids: 1, minimum V/J-germline ID>= .8.
- L1 north cluster = L1-12-B, L2 north cluster = L2-8-A, L3 north cluster = L3-8-A, maximum number of PTMs=6, maximum rare amino acids = 2, minimum V/J-germline ID >=.81.

Filters for the CMAB0-based designs included:

- H1 north cluster = H1-13-A, H2 north cluster = H2-10-A, H3 length = 10, maximum number of PTMS: 12, maximum rare amino acids: 0, minimum V/J-germline ID>= .86.
- L1 north cluster = L1-16,17-A, L2 north cluster = L2-8-A, L3 north cluster = L3-9,10-A, maximum number of PTMS=7, maximum rare amino acids = 0, V/J-germline ID>=.80.

Scaffold-CDR combinations were ranked according to Rosetta Energy, the developability metrics from Therapeutic Antibody Profiling, and sampled to maximize FR diversity as described previously.

### Expression and purification of recombinant monoclonal antibodies

The variable heavy and light chain sequence of anti-SARS CoV2 antibodies S309, and CMAB0 ^22^ and variants were cloned into the full length IgG1 expression vectors pcDNA3.3 HC and pcDNA3.3 LC (ATUM). The recombinant antibodies were transiently expressed in both Expi CHO and Expi293 cells according to manufacturer’s protocol (Invitrogen). The supernatants from 1mL transient transfections of the antibodies were purified using the AssayMAP BRAVO platform with 5mL Protein A cartridges (Agilent Technologies). Lager scale transient transfections were purified on the Akta FPLC system using 1mL HiTrap MabSelect PrismA™ affinity chromatography resin (Cytiva). The purified recombinant monoclonal antibodies were stored in 1x phosphate buffered saline at 4°C until use. Specific site directed mutations on the S309 and CMAB0 antibody sequence was done using the Quick-change site directed mutagenesis kit II (Agilent technologies).

### Screening of Expressed Recombinant Antibodies Using Enzyme Linked Immunosorbent Assay (ELISA)

The antibodies purified from a 1mL transient transfection was tested for binding against BA.1 RBD (Acro# SPD C522e) protein on an ELISA. Briefly, 2ug/mL of SARS CoV2 BA.1 RBD protein were coated on 96-well ELISA plates (Nunc Maxisorp) and left overnight at 4°C. The wells were blocked with 5% Blotto (Santa Cruz) in 1xPBST for 1hr at room temperature. Using the Opentrons OT-2 benchtop liquid handler, the purified variant recombinant antibodies based on S309 and CMAB0 were diluted to either 12 and 0.3ug/mL or 12 and 0.06ug/mL respectively and added to the plates and incubated on a rocker platform for 2hrs at room temperature. After rinsing the plates three times with 1x PBST a rabbit anti-human IgG conjugated to horseradish peroxidase (Jackson Immuno Research) was added to each well. The plates were incubated for 1hr at room temperature followed by washing with 1xPBST and addition of TMB substrate. The reaction was stopped by adding 1N sulfuric acid and the absorbance was read at 405nm.

Select CMAB and GMAB candidates were serially diluted and tested for binding against BA.1 RBD (Acro# SPD C522e) protein on an ELISA to determine apparent KD values. Briefly, 0.5ug/mL of BA.1 RBD protein were coated on 96-well ELISA plates (Nunc Maxisorp) and left overnight at 4°C. The wells were blocked with 5% Blotto (Santa Cruz) in 1xPBST for 1hr at room temperature. Using the Opentrons OT-2 benchtop liquid handler, a three-fold serial dilution of select antibodies from 9μg/mL to 0.152ng/μL was made and added to the plates and incubated on a rocker platform for 2hrs at room temperature. After rinsing the plates three times with 1x PBST a rabbit anti-human IgG conjugated to horseradish peroxidase (Jackson Immuno Research) was added to each well. The plates were incubated for 1hr at room temperature followed by washing with 1xPBST and addition of TMB substrate. The reaction was stopped by adding 1N sulfuric acid and the absorbance was read at 405nm.

### Affinity determination using Octet (biolayer interferometry)

The affinity of the antibodies to BA.1 RBD (Acro# SPD C522e), BA.2 RBD (Acro# SPD-C522g), BA.4/5 RBD (Acro# SPD-C522r), and XBB.1 RBD (Acro# SPD C5241) was determined using the Octet. The Pro A sensors were presoaked in 20mM HEPES with 150mM NaCl and 0.05% tween 20, pH 7.4 or assay buffer and then loaded with 0.5ug/mL of select recombinant monoclonal antibody. A twofold dilution of the different RBDs from 60nM to 1.875nM in assay buffer was made and the antibody coated Pro A sensors were then incubated in the various dilutions followed by dissociation in assay buffer. The KD values were calculated using the global fit method on the Octet Red96(Sartorius) instrument.

### Pseudovirus neutralization assay

For the pseudovirus neutralization assay, HEK 293T cells expressing ACE2 and TMPRSS2 were purchased from Genecopoeia (Catalog # SL222). The pseudovirus particles used in the study were procured from eEnzyme (catalog #s SCV2-PsV-Omicron, and SCV2-PsV-OmiBA2). The HEK 293T cells were maintained in DMEM (Corning Catalog# 10-013-CV) containing 10% FBS (Gibco Catalog # A38401-01) containing selection antibiotics hygromycin and puromycin as per the manufacturer’s protocol. The test antibodies were incubated with the pseudovirus particles for 1h at 37°C. Afterwards, HEK 293T cells expressing hACE2 and TMPRSS2 were incubated with the antibody/pseudovirus mixture and incubated for 48h. After 48hours incubation, the luciferase activity of SARS-CoV-2 pseudovirus infected HEK 293T cells were determined by luciferase reporter assay kit (eEnzyme, Catalog # CA-L165-10). e. The relative luciferase activity (%) was calculated as follows:

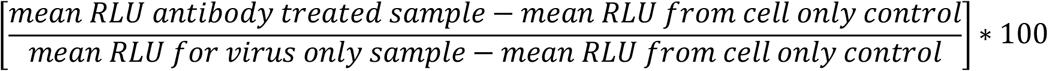

Results of neutralization assays were plotted by normalization to samples where no antibody was used, and the half-maximal inhibitory concentration (IC50) was calculated using 4-parameter non-linear regression using GraphPad Prism. Each experiment was run in duplicate.

### Analysis of Select candidates by Size Exclusion Chromatography (SEC) and Thermal Shift Assay

Briefly, the purity of the recombinant monoclonal antibody samples was studied using the AdvanceBio, 300A, 2.7um, 4.6 × 300mm (Agilent, P/N: PL1580-5301) SEC column connected to Agilent 1260 Bioinert Infinity Quaternary Pump System (Pump serial# DEAGH00678). 1X Phosphate Buffered Saline at pH 7.4 and filtered with a 0.2um membrane filter unit was used as the mobile phase. After calibrating and verifying the performance of the column using the LC Bio-standard (Agilent, P/N : 5190-9417), the recombinant antibody samples were diluted to 2mg/ml in 1XPBS and loaded onto a pre-conditioned column at a flow rate of 0.35ml/min. The samples were monitored by ultraviolet (UV) absorbance at a wavelength of 280nm with 4nm bandwidth. Using Agilent OpenLab software, based on the retention times, the Monomeric, Aggregate and Low molecular weight peaks were identified. The peak area percentage represents the relative concentration of the monomer, aggregate and fragments.

For the Thermal Shift Assay, select recombinant monoclonal antibody samples were diluted to 1.05 mg/mL in 1X Phosphate Buffered Saline at pH 7.4 and mixed with 20X SYPRO Orange Fluorescent Dye (Sigma Cat# S5692, 50uL) in a 96-well plate. The sealed plate was pulse centrifuged for up to 1000rpm to ensure there are no bubbles in the samples and run on the QuantStudio 3 device (Applied Biosystems) and the data was analyzed using their Protein Thermal Shift software.

## Notes

### Competing Interest Statement

The authors have declared no competing interest.

