## Supporting Online Information for "Machine Learning-Guided Antibody Engineering That Leverages Domain Knowledge To Overcome The Small Data Problem"

**Structural models only partly explain observed affinity enhancement of our lead mAbs.**

The analysis of the contacts between antibody and antigen across different Omicron subvariants RBDs were performed using Rosetta-modeled structural complexes (**Supplementary Methods**). These contacts are described in detail in **Supplementary Table S4**. In the case of CMAB283, increased contacts in the CMAB283 H3 mutation H99W and the epitope, including potential hydrophobic interactions with epitope residues 412P and 429F, were observed. On the light chain, there are no significant interactions in CMAB283 relative to CMAB0, despite point mutation data indicating significantly better binding for mutations such as L93K which cannot be explained by the structural model.

Both GMAB156 and S309 had variable binding across Omicron subvariants, with only GMAB156 maintaining sub nanomolar KD for all subvariants. The affinity differences between GMAB156 and S309 on the BA.1, BA.2, and BA.4 RBDs respectively may be explained by the electrostatic interactions between the GMAB156 H1 mutation 31K and 339D on the RBD and the L3 mutation T93E and RBD 346R, with additional VDW contacts between L2 mutation S52Y and 440K and polar contacts between H3 mutation A100S and RBD 340E also playing a role. In XBB.1, the D339H mutation results in significantly more polar contacts for S309 H31S, improving its binding. The salt bridge between GMAB156 H31K and 339D is replaced by a pi-cation interaction between H31K and 339H and GMAB156 maintains its sub nanomolar binding. While some of the mutations selected by our platform are rationalizable using structure-based reasoning, the platform may also discover mutations that are not obvious *a priori* from static structure analysis.

**Supplementary Figures**

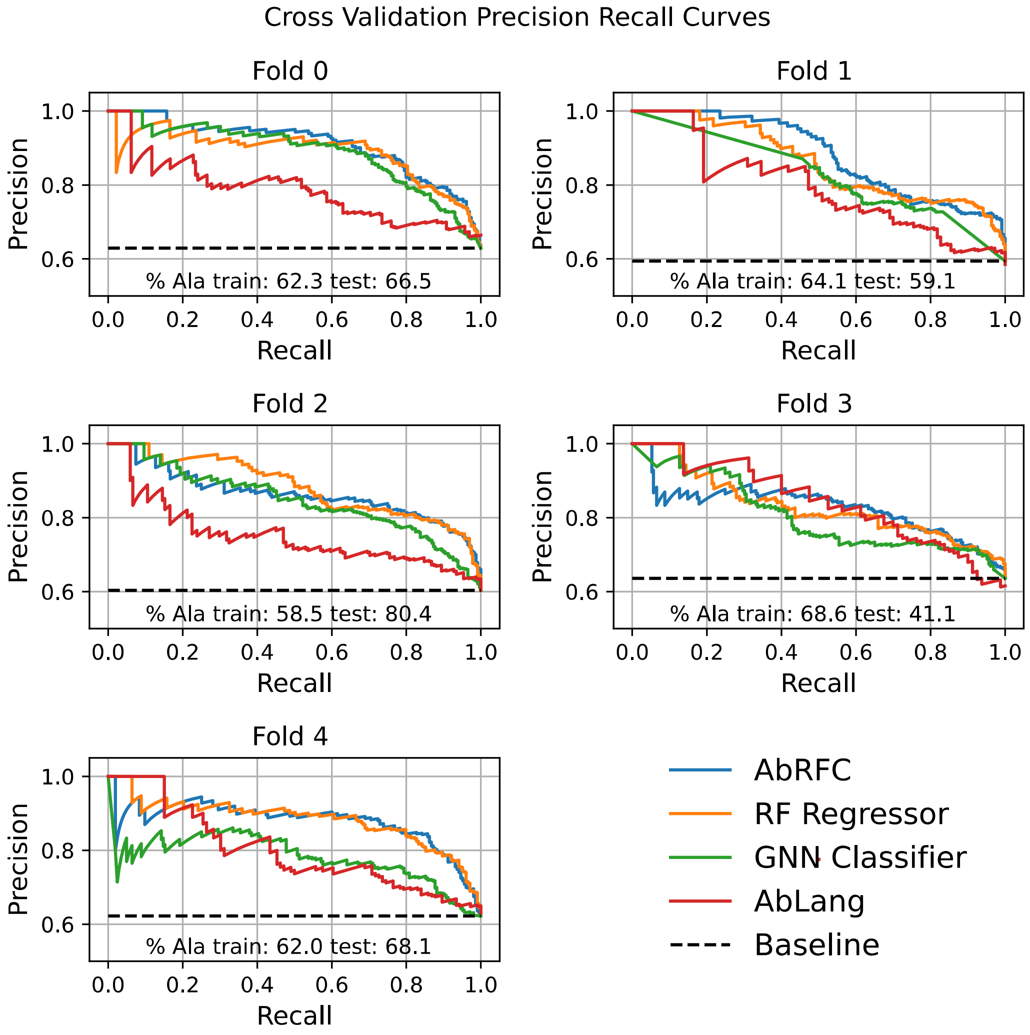


**Figure S1.** Cross-validation performance of the various models. The cutoff used (see methods) results in a random precision of ~.6. All structure-based models show relatively similar performance above baseline, while ABLANG performs surprisingly well given that it uses no epitope information and is not trained for this task.

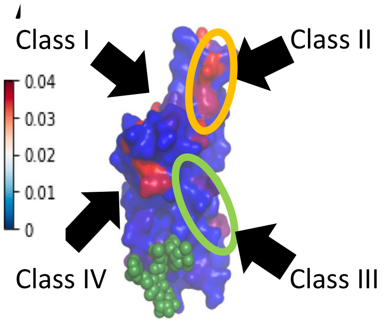


| Structure types | CoV-AbDab mAbs in Clonotype | Total CoV-AbDab mAbs | OAS Individuals | OAS Sequences | Avg Epitope Entropy |
|---|---|---|---|---|---|
| 31 | 231 | 251 | 67 | 13835 | 0.1 |
| 11 | 28 | 64 | 25 | 1249 | 0.1 |
| 2 | 4 | 24 | 7 | 65 | 0.06 |
| 5 | 13 | 22 | 22 | 543 | 0.03 |

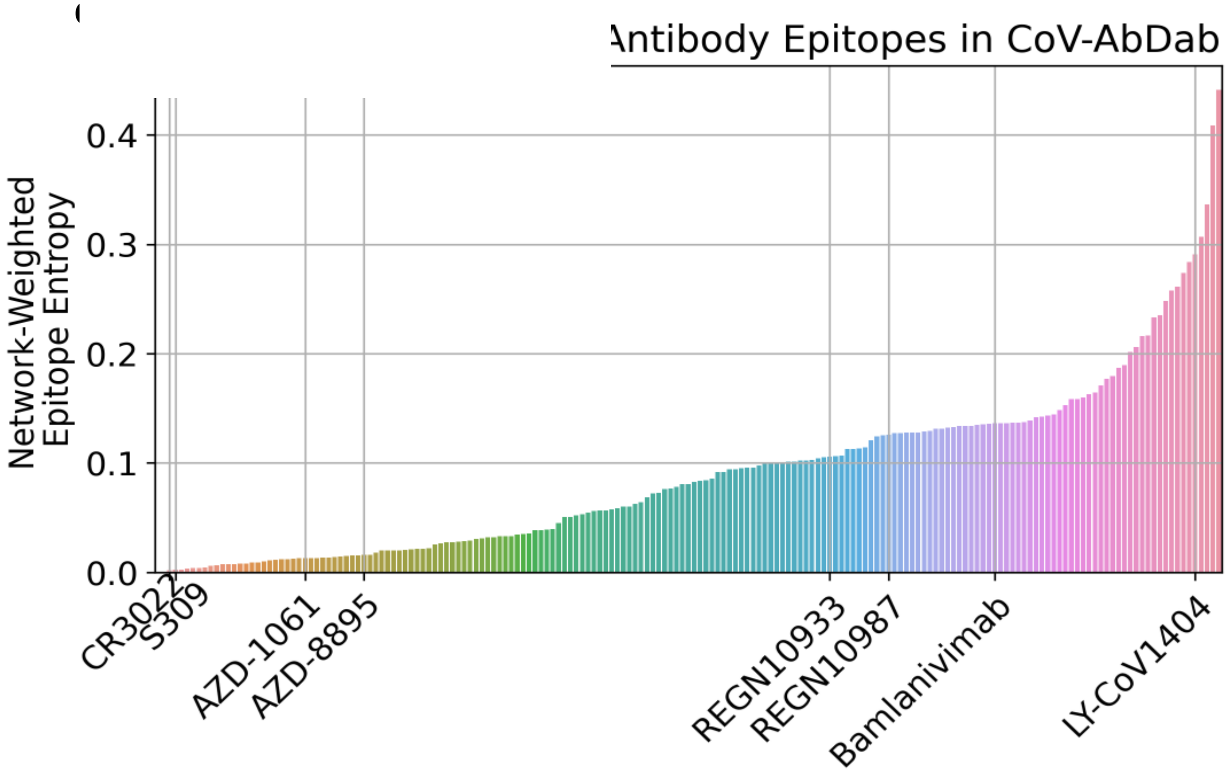


**Figure S2**. **Target epitope selection on SARS-COV-2 spike protein. A.** The residue-level entropy in the context of sequence evolution (calculated from GISAID sequences) was used to map the entropy profiles of different epitopes onto the three-dimensional structure of the RBD (shown as surface rendering in purple). The Class III epitope region (encircled in green) encompassing a surface glycosylation (colored green) targeted by antibodies such as S309 and Class IV epitope region targeted by antibodies such as CR3022 have a substantially lower entropy than Class II epitope region targeted by antibodies such as Bamlanivimab (encircled in orange). **B.** Analysis of structures available in CoV-AbDab showed clonotypes. When additional Abs from CoV-AbDab and OAS were mapped to these clonotypes, a bias for ACE2 blocking class I and class II Abs was observed both in the CoV-AbDab subset and the OAS subset. These Ab classes also have, on average, the highest epitope entropies. **C.** Unique class III clonotype S309 and class IV clonotype CR3022 target extremely low-entropy epitopes.

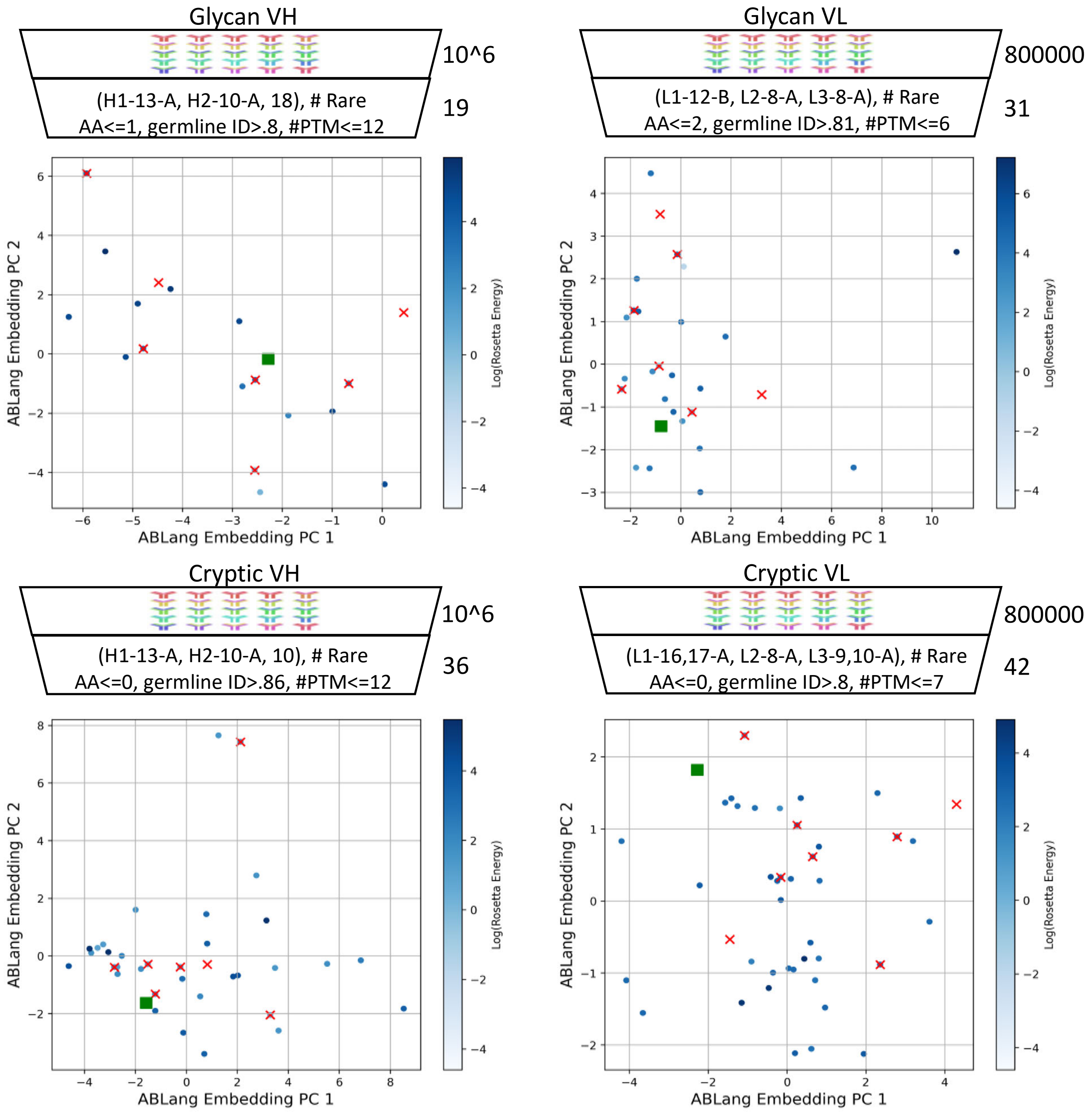


**Figure S3.** PCA analysis of the ABLang embeddings of candidate and selected scaffolds. Scaffolds before filtering are blue dots, after filtering are red crosses, and selected for the final lead candidates are green squares. The spread of the filtered scaffolds shows the emphasis of diversity in the scaffold selection process.

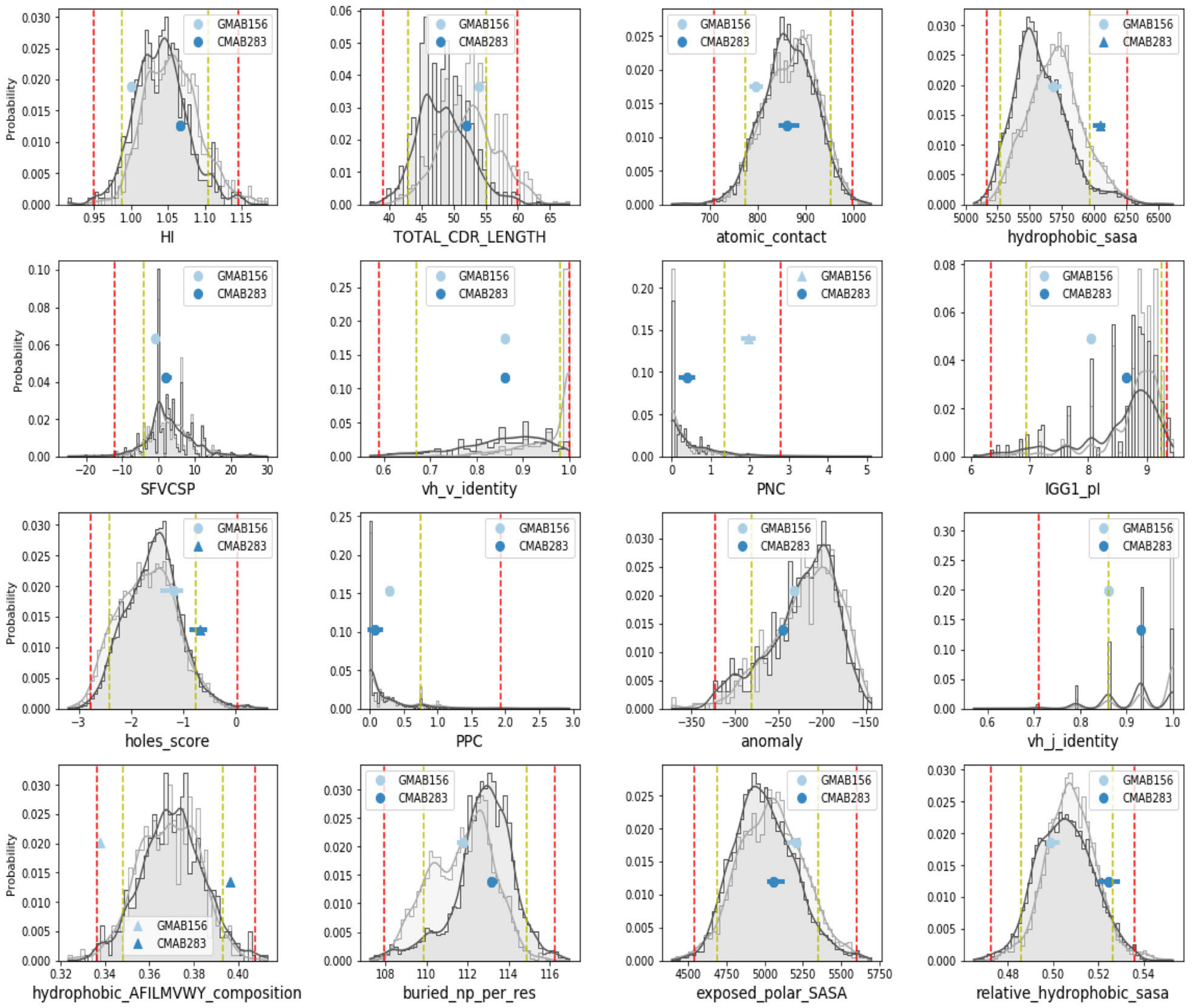


**Figure S4.** *In Silico* developability profile of the lead candidates. The dark gray distribution is the distribution of ~500 CSTs across these *in silico* metrics, while the light gray distribution is the distribution of 500 structure-matched antibodies from OAS. The yellow and red lines represent the outside 10% and 5% of the distribution for each metric. The metrics are meant to capture three orthogonal subsets of *in silico* metrics that correlate with the developability of an antibody: structure independent (sequence level) metrics, structural metrics related to the surface of the antibody, and structural metrics related to the buried core including the VH/VL interface (packing level). **Sequence level metrics:** HI: hydrophobicity index, TOTAL_CDR_LENGTH, IGG1_pI (pI of the antibody in the context of an IGG1 FC), anomaly: sum of log probabilities of observing each amino acid at each position in human OAS, vh_v_identity, vh_j_identity, hydrophobic_AFILMVWY_composition. **Structure level surface metrics:** hydrophobic_sasa, SFVCSP (TAP), PNC (TAP), PPC (TAP), exposed_polar_sasa, relative_hydrophobic_sasa. **Structure level packing metrics:** atomic_contact: sidechain carbon-carbon counts, holes_score: relative number of voids in the protein core (https://www.rosettacommons.org/docs/latest/scripting_documentation/RosettaScripts/Filters/HolesFilter) , buried_np_per_res: buried non-polar SASA per residue.

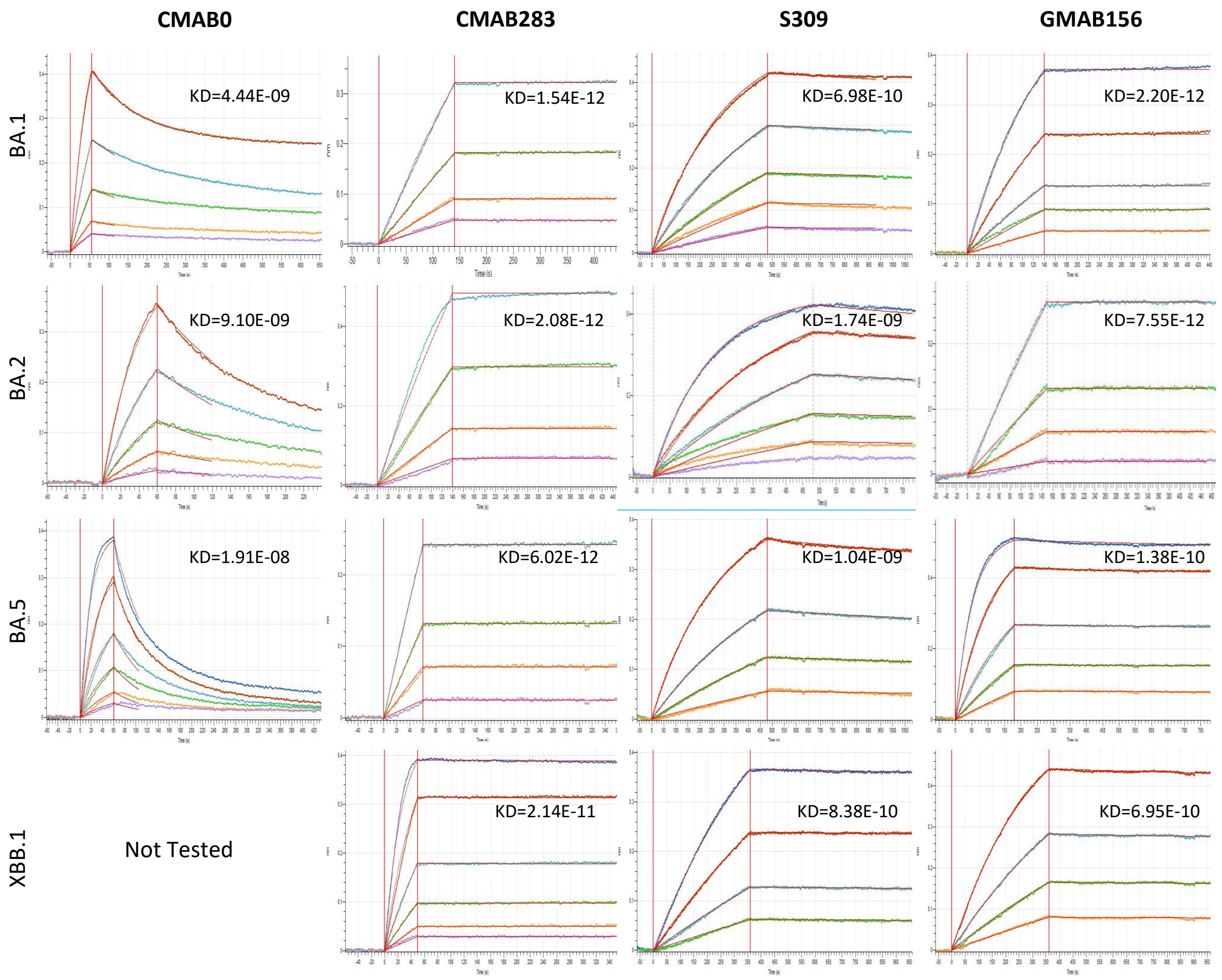


**Figure S5.** Octet BLI curves for the templates and lead candidates against the VOC sublineages of BA.1. Candidates were designed against BA.1 but retain their sub-nanomolar binding against BA.2, BA.5, and XBB.1.

### Supplementary Tables

| Feature Name | Description | Source |
|---|---|---|
| idSASA_fraction_1 | dSASA_fraction (Mutant Position Neighbors) | PyRosetta |
| idhSASA_sc_0 | dhSASA_sc (Mutant Position) | PyRosetta |
| idSASA_0 | dSASA (Mutant Position) | PyRosetta |
| dE | Rosetta ref2015 scorefxn (Full Complex) | PyRosetta |
| iinterface_residues_1 | # interface residues (Mutant Position Neighbors) | PyRosetta |

| | | |
|---|---|---|
| idSASA_fraction_0 | dSASA_fraction (Mutant Position) | PyRosetta |
| idSASA_sc_0 | dSASA_sc (Mutant Position) | PyRosetta |
| fa_rep_1 | fa_rep (Mutant Position Neighbors) | PyRosetta |
| hbond_bb_sc_0 | hbond_bb_sc (Mutant Position) | PyRosetta |
| fa_atr_1 | fa_atr (Mutant Position Neighbors) | PyRosetta |
| iinterface_hbonds | # hbonds (Interface Residues) | PyRosetta |
| isc_value | Shape Complementarity (Full Complex) | PyRosetta |
| idelta_unsat_hbonds | delta_unsat_hbonds (Interface Residues) | PyRosetta |
| fa_elec_0 | fa_elec (Mutant Position) | PyRosetta |
| sin_res | Residue SIN Value (Mutant Position) | SIN |
| sin_norm | SIN Normalization Constant (Full Complex) | SIN |
| lk_ball_wtd_1 | lk_ball_wtd (Mutant Position Neighbors) | PyRosetta |
| fa_atr_0 | fa_atr (Mutant Position) | PyRosetta |
| total_score_0 | total residue score (Mutant Position) | PyRosetta |
| fa_sol_0 | fa_sol (Mutant Position) | PyRosetta |
| hbond_bb_sc_1 | hbond_bb_sc (Mutant Position Neighbors) | PyRosetta |
| total_score_1 | total residue score (Mutant Position Neighbors) | PyRosetta |
| hbond_sc_0 | hbond_sc (Mutant Position) | PyRosetta |
| idhSASA_sc_1 | dhSASA_sc (Mutant Position Neighbors) | PyRosetta |
| fa_elec_1 | fa_elec (Mutant Position Neighbors) | PyRosetta |
| idG_1 | ddG (Mutant Position Neighbors) | PyRosetta |
| fa_sol_1 | fa_sol (Mutant Position Neighbors) | PyRosetta |
| aif_score | Residue AIF Score (Mutant Position) | AIF |
| sin_if | SIN (Interface Residues) | SIN |
| idhSASA_1 | dhSASA (Mutant Position Neighbors) | PyRosetta |
| fa_rep_0 | fa_rep (Mutant Position) | PyRosetta |

**Table S1.** Description of the features used in the AbRFC model. Each feature is either calculated at the level of the complex, the interface, the mutant residue, or the neighbors of the mutant residue (see **Description** column). Features come from 3 sources – the AIF calculation, the SIN calculation, or features described in Pyrosetta. The calculation of a node's neighbors is defined in the **Supplementary Methods.**

**Table S2** (separate excel sheet). The list of filtered scaffolds considered for porting the affinity enhanced loops. Along with the sequence of the scaffold, 5 *in silico* metrics used during the scaffold selection process are reported: $\Delta E$ is the difference in total Rosetta energy of porting the loop onto the scaffold relative to keeping the loop on the template scaffold. pKA Score – sum of pKA values based on sequence. PSH, PPC, PNC, SFVCSP scores are calculated using the formula from the TAP web server.

| Measurement | GMAB156 | CMAB283 |
|---|---|---|
| **Tm (°C)** | 65.5 | 70.9 |
| **SEC Monomer (Peak Area %)** | 97.5 | 99.3 |

**Table S3**. Experimental developability characterization of the round 2 mAbs shows that they Tm and SEC profiles that are within the range for therapeutic mAbs.

**Supplementary Methods**

__RandomForestClassifier__

**Neighbor calculation for features:**
Some features are calculated over node neighbors (**Table S1**). In these cases, the neighbors were defined as follows:

$$\mathrm{NBRS}(res_i) = res_j\ s.t\ \max\left(E\left[res_i, res_j\right]\right) > c$$

Where $\max(E[i,j])$ is the maximum absolute value of the considered Pyrosetta Energy terms (total_score, fa_atr, fa_rep, fa_sol, fa_intra_rep, fa_intra_sol_xover4, lk_ball_wtd, fa_elec, pro_close', hbond_sr_bb, hbond_lr_bb, hbond_bb_sc, hbond_sc) between residues i and j. Second order neighbors were considered by applying the above formula recursively, with c=.05 for first order neighbors and c=.1 for second order neighbors.

**Parameters:**
```
`{'bootstrap': True,
  'class_weight': None,
  'criterion': 'entropy',
  'max_depth': 50,
  'max_features': 'auto',
  'max_leaf_nodes': None,
  'min_impurity_decrease': 0.0,
  'min_samples_leaf': 10,
  'min_samples_split': 10,
  'min_weight_fraction_leaf': 0.0,
  'n_estimators': 1000,
  'n_jobs': None,
  'oob_score': False,
```

'random_state': None,
'verbose': 0,
'warm_start': False}`

**RandomForestRegressor**

**Parameters:**

```
{'bootstrap': True,
 'max_depth': 50,
 'max_features': 'sqrt',
 'max_leaf_nodes': None,
 'min_samples_leaf': 10,
 'min_samples_split': 10,
 'min_weight_fraction_leaf': 0.0,
 'n_estimators': 1000,
 'n_jobs': None,
 'oob_score': False,
 'random_state': None,
 'verbose': 0,
 'warm_start': False}
```

**Contact Analysis**

**get_static_contacts.py parameters**

```
"--output", out_file,
"--itypes", "all",
"--sb_cutoff_dist", '5.0', #cutoff for distance between anion and cation atoms [default = 4.0 angstroms]
"--pc_cutoff_dist", '7', #cutoff for distance between cation and centroid of aromatic ring [default = 6.0 angstroms]
"--pc_cutoff_ang", '80', #cutoff for angle between normal vector projecting from aromatic plane and vector from aromatic center to cation atom [default = 60 degrees]
"--ps_cutoff_dist",'7',  #cutoff for distance between centroids of two aromatic rings [default = 7.0 angstroms]
"--ps_cutoff_ang",'50',  #cutoff for angle between the normal vectors projecting from each aromatic plane [default = 30 degrees]
"--ps_psi_ang",'60',     #cutoff for angle between normal vector projecting from aromatic plane 1 and vector between the two aromatic centroids [default = 45 degrees]
"--ts_cutoff_dist",'6',  #cutoff for distance between centroids of two aromatic rings [default = 5.0 angstroms]
"--ts_cutoff_ang",'45',  #cutoff for angle between the normal vectors projecting from each aromatic plane minus 90 degrees [default = 30 degrees]
```

"--ts_psi_ang",'60',     #cutoff for angle between normal vector projecting from aromatic plane 1 and vector between the two aromatic centroids [default = 45 degrees]
"--hbond_cutoff_dist",'6',#cutoff for distance between donor and acceptor atoms [default = 3.5 angstroms]
"--hbond_cutoff_ang", '180',#cutoff for angle between donor hydrogen acceptor [default = 180 degrees]
"--hbond_res_diff",'1',#minimum residue distance for which to consider computing hbond interactions [default = 1]
"--vdw_epsilon",'3',   #amount of padding for calculating vanderwaals contacts [default = 0.5 angstroms]
"--vdw_res_diff",'2'], #minimum residue distance for which to consider computing vdw interactions [default = 2]]

### Epitope Identification

***Clonotype Identification:*** Clonotypes were first identified on the 186 SARS-COV-2 RBD targeting mAbs with structures available in SAbDab at the time of the analysis. Distance between 2 mAbs was computed as:

$$\mathrm{D_{ij}} = 1 - \min(\mathrm{ID}(\mathrm{VH_1}, \mathrm{VH_2}), \mathrm{ID}(\mathrm{VL_1}, \mathrm{VL_2}))$$

Where $\mathrm{ID}(\mathrm{SEQ_1}, \mathrm{SEQ_2}) = \frac{\sum_{pos=1}^{N} \mathrm{SEQ_1(pos)=SEQ_2(pos)},}{\max\,(\mathrm{len(SEQ_1)},\mathrm{len(SEQ_2)})}$By iteratively computing clonotypes using agglomerative clustering with the complete linkage method and a threshold cutoff *t* such that no two sequences in the same cluster will have $D_{ij} > t$, it was found that *t*=.12 reliably clustered only those mAbs targeting highly overlapping epitopes (**Table S1**). This threshold was then used to identify clonotypes across all sequences downloaded CoV-AbDab.

***Epitope and Class Definition:*** Epitope-paratope contacts were identified as any contacts with heavy atom distance <5A. To assign RBD residues to epitope classes, the mAb assignments from the (Barnes et al) paper were used to identify all residues that were in the epitope of any mAb of a certain class. For example, C102 is in Barnes class I and 486 is in the C102 epitope, therefore 486 is a class I epitope residue. Note that a residue may belong to multiple Barnes classes (486 is a class I and class II residue). MAbs not classified by Barnes were assigned to classes if 40% of their epitope residues were in a Barnes class.

***RBD Residue Entropy Calculation:*** Spike protein sequences were downloaded from GISAID and filtered to only include human full length spike sequences. Unique sequences were aligned using the augur component of the NextStrain python package. The Sequence Entropy at each position was calculated as follows:

$$f_{R,aa} = \frac{\#\,Sequences(AA_R = aa)}{Total\ \#\ Sequences}$$

$$\text{Sequence Entropy(R)} = -\sum_{\text{aa}} f_{R,AA} \log f_{R,AA}$$

Here $f_{R,aa}$ is the frequency of $aa$ appearing at $R$ in the dataset and $\text{Sequence Entropy(R)}$ is the Shannon Entropy at R given the $f_{R,aa}$ distribution.

***Network Weighted Entropy Calculation:*** The total entropy of a mAb epitope (used in figure 2C) was calculated as follows:

$$\text{Entropy}(Epitope) = \sum_{R \in Epitope} (\text{Sequence Entropy}(R) * \sum_{P \in Paratope} \text{Network}(R, P))$$

Where R is the epitope residue, P is a paratope residue, Network(R,P) was calculated using Significant Interaction Network between R and P as described previously [32] and Sequence Entropy of R was calculated using the GISAID data.

### Structural Analysis of Mutations

Models of CMAB283 and GMAB156 were obtained by applying the mutations to the 7BEP and 7LOP-based models described above. Contacts were defined using the get_static_contacts.py script from https://github.com/getcontacts/getcontacts. Flags for computing the various interactions are given in the supplement. Only interactions between the antibody and the RBD were considered, and a unique interaction was defined by (interaction_type, epitope residue number, paratope chain, paratope residue number).
